# Juvenile Shank3 KO mice adopt distinct hunting strategies during prey capture learning

**DOI:** 10.1101/2022.06.13.495982

**Authors:** Chelsea Groves Kuhnle, Micaela Grimes, Victor Manuel Suárez Casanova, Gina G. Turrigiano, Stephen D. Van Hooser

## Abstract

Mice are opportunistic omnivores that readily learn to hunt and eat insects such as crickets. The details of how mice learn these behaviors and how these behaviors may differ in strains with altered neuroplasticity are unclear. We quantified the behavior of juvenile wild type and Shank3 knockout mice as they learned to hunt crickets during the critical period for ocular dominance plasticity. This stage involves heightened cortical plasticity including homeostatic synaptic scaling, which requires Shank3, a glutamatergic synaptic protein that, when mutated, produces Phelan-McDermid syndrome and is often comorbid with autism spectrum disorder (ASD). Both strains showed interest in examining live and dead crickets and learned to hunt. Shank 3 knockout mice took longer to become proficient, and, after 5 days, did not achieve the efficiency of wild type mice in either time-to-capture or distance-to-capture. Shank3 knockout mice also exhibited different characteristics when pursuing crickets that defied explanation as a simple motor deficit. Although both genotypes moved at the same average speed when approaching a cricket, Shank3 KO mice paused more often during approaches, did not begin final accelerations toward crickets as early, and did not close the distance gap to the cricket as quickly as wild type mice. These differences in Shank3 KO mice are reminiscent of some behavioral characteristics of individuals with ASD as they perform complex tasks, such as slower action initiation and completion. This paradigm will be useful for exploring the neural circuit mechanisms that underlie these learning and performance differences in monogenic ASD rodent models.

## Introduction

Small rodents such as rats and mice are prey for many larger animals, but they are also opportunistic hunters that learn to pursue, capture and eat small insects such as crickets (Berry and Bronson, 1992; Hoy et al., 2016; Galvin et al., 2021). Cricket hunting by mice has been studied carefully in the laboratory. Adult mice use their hands and mouths to grab crickets, and, over successive days, rapidly decrease the amount of time needed to capture and eat crickets in an enclosure (Hoy et al., 2016; Galvin et al., 2021). While mice can perform this task using auditory or somatosensory information, vision is a key sense that is used under normal circumstances (Hoy et al., 2016). When an adult mouse pursues a cricket, the mouse orients its head directly toward the cricket and maintains a lowered head posture (Michaiel et al., 2020) that places the cricket in the portion of the visual field of both retinas where the density of alpha-ON ganglion cells is highest and the optic flow is the smallest (Holmgren et al., 2021). Naïve mice are attracted to the appearance of a small cricket-sized spot, and mice with experience hunting crickets are even more attracted to these stimuli (Procacci et al., 2020).

The critical period for ocular dominance plasticity is a time of great change for the visual system (Wiesel and Hubel, 1965; Gordon and Stryker, 1996; Wang et al., 2010; Chang et al., 2020), as both Hebbian and homeostatic mechanisms are employed to ensure proper response gain and the alignment of signals from the two eyes (Heynen et al., 2003; Frenkel and Bear, 2004; Kaneko et al., 2008; Maffei and Turrigiano, 2008; Lambo and Turrigiano, 2013). These plasticity processes may be enhanced by the performance of complex ethological tasks, and we were therefore interested in characterizing in detail how critical period mice (aged postnatal day 28-30) learn to perform this vision-dependent behavior, and determine whether this learning is altered in mice with impaired visual system plasticity. Loss of Shank3, a major scaffolding protein at glutamatergic synapses (Monteiro and Feng, 2017), is strongly associated with Phelan-McDermid syndrome and autism-spectrum disorders (ASDs), (Leblond et al., 2014; Costales and Kolevzon, 2015), and is required for normal hippocampal LTP (Bozdagi et al., 2010; Song et al., 2019), and the expression of homeostatic and ocular dominance plasticity within V1 ((Tatavarty et al., 2020)Tatavarty et al., 2020; Wu et al., 2022).

We were also interested in studying the process of learning a complex behavior in a Shank3 ASD mouse model for more general reasons. Humans with autism spectrum disorder (ASD) require more time to plan and execute goal-directed movements, exhibit more temporal and spatial variability during initial movement, and move at slower speeds ((Glazebrook et al., 2006; Longuet et al., 2012). In addition, ASD individuals demonstrate less flexible responses in performing goal-directed actions (Alvares et al., 2016), and diminished audio-visual multisensory integration (Stevenson et al., 2014; Brandwein et al., 2015; Feldman et al., 2018). Shank3 KO mice display impaired multisensory integration, hyperreactivity to tactile sensory input, altered audio-tactile responses, and weakened auditory responses (Gogolla et al., 2014; Engineer et al., 2018; Chen et al., 2020). A detailed understanding of how autism-associated mutations contribute to difficulty in learning ethologically relevant tasks is lacking.

To examine these questions we allowed critical period Shank3 knockout mice and their wild type littermates to hunt crickets over a period of five days. Both strains learned to hunt crickets, but Shank3 KO mice were initially less likely to recognize crickets as prey and initiate hunting, and took longer to acquire the behavior. By day five, Shank3 KO mice captured and consumed as many crickets as wild type animals, and were only a little slower in overall time-to-capture. However, there were two major differences in the approach phase of the behavior of Shank3 KO and wild type animals. First, Shank3 KO mice paused much more frequently during their approaches to crickets. Second, while Shank3 KO and wild type mice exhibited the same average and peak speeds during approaches, wild type mice showed much greater modulation of their speed, increasing their speed progressively as they neared a cricket in a way that Shank3 KO mice did not. Therefore, just as neurotypical individuals and autism spectrum disorder individuals exhibit different behavioral strategies when performing goal-directed tasks ((Willshaw et al., 1969; Glazebrook et al., 2006; Alvares et al., 2016), wild type and Shank3 KO mice developed cricket hunting behaviors with distinct features. The circuit mechanisms underlying these differences can be explored in future experiments.

## Methods

### Experimental Model and Subject Details

All procedures were approved by the Brandeis University Institutional Animal Care and Use Committee, and conformed to the National Institutes of Health Guide for the Care and Use of Laboratory Animals. For all experiments, mice of both sexes were used. No differences were noted between males and females and data were combined. We used the homozygous Shank3B knockout mouse introduced by Peça et al. (2011), obtaining founder mice from Jackson labs (Stock No: 017688). Genotyping was done based on primers as previously described (Peca et al., 2011). All animals began habituation between the ages of p26-p28 and data were combined.

### Live Cricket hunting

Littermate pairs of animals were moved to the behavioral testing room two days before testing and housed as previously described (Hengen et al., 2013). Cedar chip bedding, standard chow, water, huts and several toys were provided in the enclosure. A twelve-hour circadian cycle was maintained, with dark hours between 7:30 pm and 7:30 am. Animals were weighed each day and monitored for health indicators. During the second night of habituation, mice were deprived of standard chow for no more than 16 hours total and five decapitated crickets were left in the enclosure. Ad libitum access to water continued during food deprivation. On the morning of the third day, testing began. The enclosure in which animals were housed was used additionally as the arena for hunting. Animals were moved to a carrier after weighing, while the arena was prepared. Bedding was disposed of and Fisherbrand™ Absorbent Underpads were taped down both on the underside and at the edges of the arena. On each corner of the arena, a cricket dispenser was placed and loaded so that an Arduino motor system could be used to robotically dispense crickets. Dispensers on opposite corners of the arena were programmed to rotate simultaneously so that animals would not be able to predict which dispenser the cricket was dropping from. Animals were first habituated to the arena for several minutes before a cricket was released. Behavior was recorded using a Logitech C920x HD Pro Webcam and Synapse (TDT) software. The duration of each hunting session was either three hours, or up to 6 crickets per session, whichever was shorter. One hunting session per day, per animal, was performed. Once both animals completed the hunting task, the absorbent padding was removed from the arena and fresh bedding was laid down along with food, huts, and toys. Water was available ad libitum throughout the experiment. After the first day of hunting, animals were food deprived without access to standard chow or dead crickets each night for a maximum of 16 hours. Animals were weighed each day and removed from the experiment if weight loss exceeded 20% of the starting weight, and continually monitored during experiments for signs of pain or distress. Five days of testing were performed per mouse. Animals were sacrificed following the fifth day of testing in accordance with the IACUC approved protocol.

### Approach and consumption during dead cricket and cereal trials

The procedure for dead cricket consumption experiments was similar to that described for live cricket hunting. Littermate pairs of mice were habituated in an enclosure, and food deprivation was performed overnight following the second day of habituation. For these tests, mice did not have access to dead crickets during food deprivation. Dead crickets used for consumption trials were decapitated the preceding night. The arena was prepared and behavior was recorded as described in the live cricket hunting tests. A mouse was placed in the arena and crickets were introduced one at a time by hand. Crickets were left in the arena for mice to consume at will. The duration of a session was three hours or after a maximum of 5-6 crickets were consumed. Mice were tested in sequence, with two dead cricket consumption trials performed per day. A single day of testing was performed for each mouse. After animals completed the dead cricket test, pieces of brightly colored sugary cereal (Froot Loops, Kellogs) were weighed and dispensed into the arena by hand one at a time. This test lasted for a maximum of one hour or until three total Froot Loops were given to the animal. Remaining Froot Loops were again weighed following consumption. Animals were sacrificed following Froot Loop tests in accordance with the IACUC approved protocol.

### Data Analysis

Videos recorded during live cricket hunting and dead cricket consumption trials were manually scored in order to determine the times of a) cricket release, b) mouse initially orienting to the cricket’s position, c) cricket capture, and d) cricket consumption. Additionally, for days one and five of the live hunting experiments, approach periods were manually scored.

Approaches were taken as periods during which the animal actively pursued the cricket, which may or may not include interceptions or end in capture. Based on manual scoring, time to capture per trial was calculated as the time between cricket release and capture. The number of approaches per trial was evaluated as the number of manually scored approaches between cricket release and capture.

DeepLabCut (DLC) (Mathis et al., 2018; Nath et al., 2019) was used to assess cricket position as well as the mouse position and orientation during trials. Still frames from hunting videos were chosen and manually labeled with cricket position and the position of the mouse body, head, nose, left ear and right ear to produce training and test data. A ResNet-50 neural network was trained on these manually labeled datasets. Following training, the performance of the network was evaluated using test data. Outlier frames were chosen and manually labeled for subsequent training iterations. Nine training iterations were performed, and the resulting network exhibited a test error of 2.0 pixels (equivalent to 0.18 cm). This network was then used to analyze the positions of mouse and cricket markers during live cricket hunting and dead cricket consumption experiments.

Outputs from DLC were analyzed using a Matlab script. Periods of under 50 frames (2.5 seconds) in which the DLC network confidence was below 98% were interpolated. Additionally, cricket positioning was improved by designating ‘jump’ frames as frames in which the decoded cricket position moved more than 10 cm between one frame and the next. ‘Jump sequences’ were considered to be time intervals consisting of multiple ‘jump’ frames with 10 frames or fewer between any individual jump frame and following jump frame. Cricket position was interpolated between the first and last position in each ‘jump sequence.’ Speed was calculated based on the change in mouse body position between one frame and the next, and smoothed by applying a sliding window average over two frames. Azimuth was calculated using cricket position, and the mouse head and nose position. Mouse-Cricket-Distance (MCD) was calculated as the distance between the mouse nose and cricket position. Interceptions were taken as frames during which the MCD fell below a threshold of 5 cm. Time immobile was calculated as time during which the mouse speed was below an immobile threshold of 1 cm/s. Distant approaches were calculated as approaches for which the MCD on the first frame of the approach was greater than a threshold value of 15 cm.

### Statistics

A Wilcoxon rank test was used for analysis in **Fig.1 D-F**, as these data corresponded to a small, discrete numeric variable (number of crickets captured or consumed) which was not normally distributed. For all other data presented as bar charts, including **Fig.1 G-J, Fig. 2, Fig. 3**, and **Fig. 4 A, B, F**, either a t-test or repeated measures ANOVA was used for statistical analysis. A value of p=0.05 was used as a threshold for statistical significance. In producing the graphs shown in **Fig. 4 G-H**, sequences of mouse speed data were identified in the second preceding and following cricket interception. A speed distribution was obtained for each frame in this sequence over the four conditions analyzed (namely naive wild type and knockout animals, and experienced wild type and knockout animals). For both naive and experienced mice, genotype speed distributions at each corresponding frame in the sequence were compared using a two-tailed t-test.

**Figure 1:**
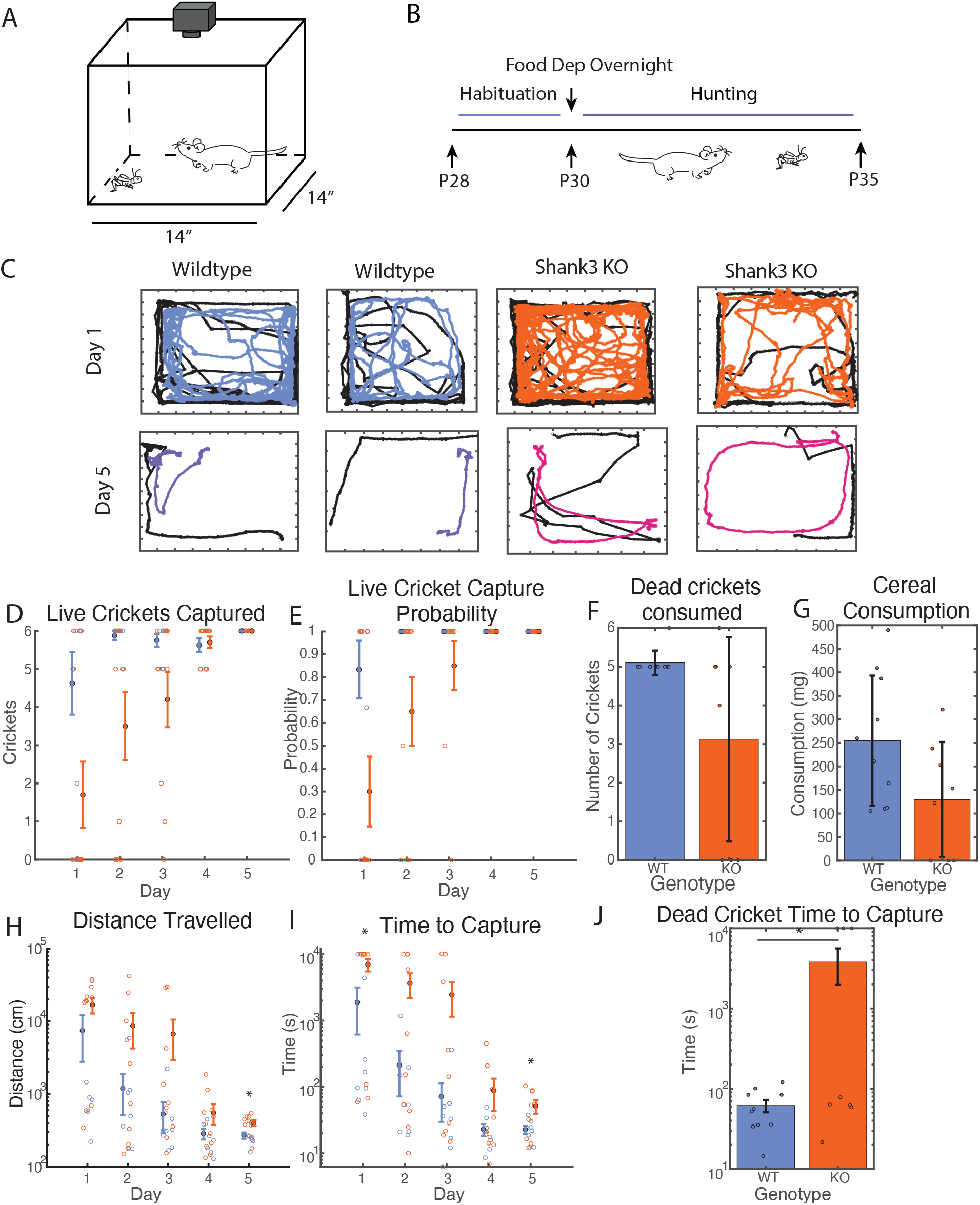
Shank3-KO animals initially consume fewer crickets than wildtype littermates and improve at hunting more slowly. (A) Schematic of recording setup for hunting sessions. Camera records behavior of mouse and cricket in a given session. (B) Experiment timeline whereby animals are habituated for two days in advance of hunting in littermate pairs. Food deprivation happens overnight the second night before beginning hunting. (C) Representative traces of mouse and cricket position during hunting trials by naive and experienced animals, in which the black trace corresponds to cricket position and the colored line corresponds to the mouse position. (D) Number of live crickets captured per session. Stats for WT vs KO, Number of crickets captured per day, Day 1 p=0. 042, Day 2 p=0. 052, Day 3 p=0. 142 (Wilcoxon). Open dots are individual observations, filled dots represent averages across multiple sessions and error bars represent standard error. (E) Fraction of cricket trials concluding in successful capture per session. Stats for WT vs KO, Capture probability, Day 1 p=0. 046, Day 2 p=0. 137, Day 3 p=0. 588 (Wilcoxon). (F) Number of dead crickets consumed by naïve mice in a single control set, p=0.079 (Wilcoxon). Without training, Shank3 KO mice consume fewer dead crickets than wild type mice, suggesting there is a Shank 3KO deficit in recognizing the crickets as food or a willingness or ability to eat them. (G) Mass of sugary cereal consumed in a single control session by naïve mice, p=0.062 (t-test). (H) Median mouse distance travelled per hunting session, across multiple live cricket trials, Day 1 p=0.145, Day 2 p=0.156, Day 3 p=0.168, Day 4 p=0.205, Day 5 p=0.016 (t-test). After training, wild type mice traveled less distance to capture crickets compared with Shank3 KO mice. (I) Median time to capture per hunting session, across multiple live cricket trials. Maximum time was plotted as 10,000 seconds in the case that animals did not capture any crickets during their allotted 3 hours. Stats for WT vs KO, Time to capture, by session, Day 1 p=0.023, Day 2 p=0.055, Day 3 p=0.127, Day 4 p=0.216, Day 5 p=0.042. After training, wild type mice were slightly faster than Shank3 KO mice. (J) Time for naïve mice to capture a dead cricket in a single control set, median by session p=0.035 (t-test). Shank3 KO mice took substantially longer time to eat dead crickets than wild type mice, even when they were not moving.

**Figure 2:**
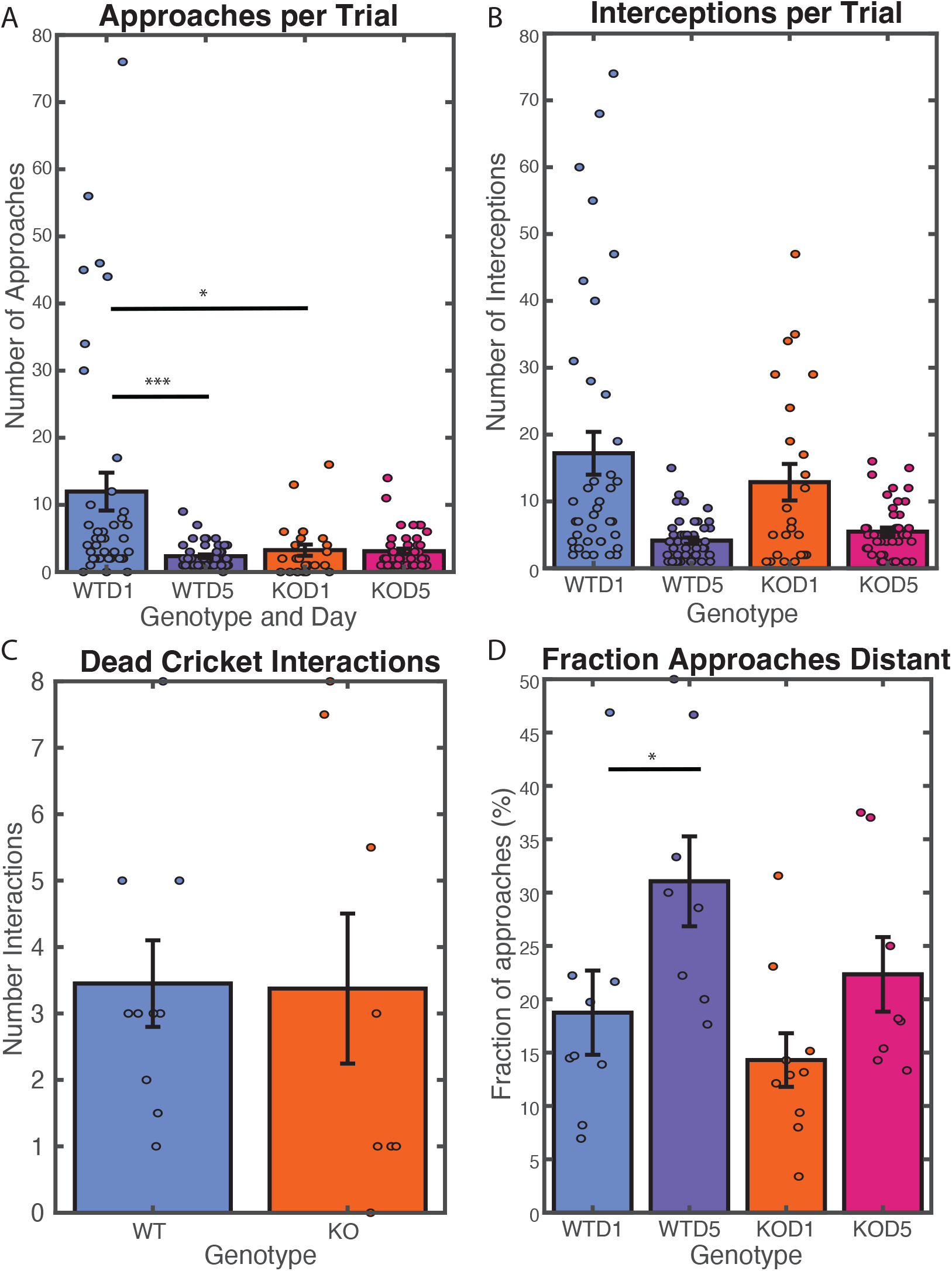
Both wild type and Shank3 KO animals gain some proficiency at cricket hunting and reduce overall number of interceptions per trial. (A) Number of approaches before cricket capture, per trial, during hunting. All stats calculated using a repeated measures ANOVA WTD1-KOD1 p=0.022, WTD5-KOD5 p=0.131, WTD1-WTD5 p=6.088e-05, KOD1-KOD5 p=0.966. (B) Number of interceptions before cricket capture, during hunting, per trial. WTD1-KOD1 p=0.352, WTD5-KOD5 p=0.365, WTD1, KOD1-KOD5 p=0.023. (C) Number of interceptions per dead cricket by naïve mice in a single control set, median by session p=0.953 (t-test). (D) Fraction of approaches for which the mouse to cricket distance exceeded 15 cm at start of approach during hunting, median per day. WTD1-KOD1 p=0.423, WTD5-KOD5 p=0.134, WTD1-WTD5 p=0.019, KOD1-KOD5 p=0.113.

**Figure 3:**
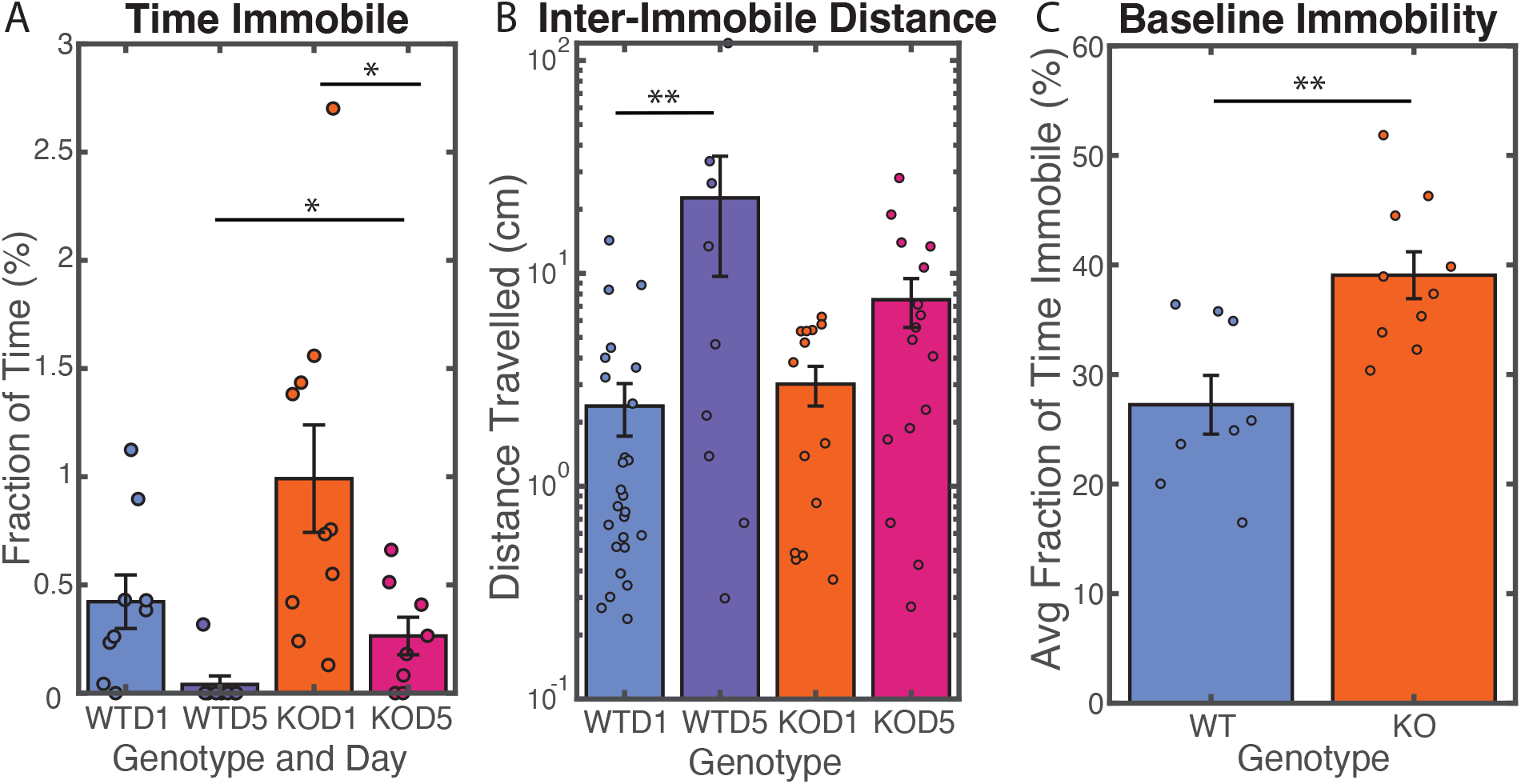
Shank3 KO mice exhibit more frequent periods of immobility than wild type mice. (A) Fraction of time immobile during approach per hunting session, averaged over all approaches. Bars show average value and error bars indicate standard error. Both genotypes spent less time immobile with training, but Shank3 KO animals continued to exhibit substantial time immobile after training whereas wild type animals exhibited very little time immobile during approaches. Calculated using repeated measure ANOVA, WTD1-KOD1 p=0.079, WTD5-KOD5 p=0.034, WTD1-WTD5 p=0.161, KOD1-KOD5 p=0.009. (B) Mean distance travelled between periods of immobility during hunting. Wild type animals traveled farther than Shank3 KO animals before pausing. Bars show average value and error bars indicate standard error. Calculated using repeated measure ANOVA, WTD1-KOD1 p=0.959, WTD5-KOD5 p=0.189, WTD1-WTD5 p=0.028, KOD1-KOD5 p=0.443. (C) Average fraction of time immobile in the absence of appetitive stimulus, by session. Time immobile calculated using two-sample t-test, p=0.003. Shank3 KO mice spent more time immobile when no cricket was present.

**Figure 4:**
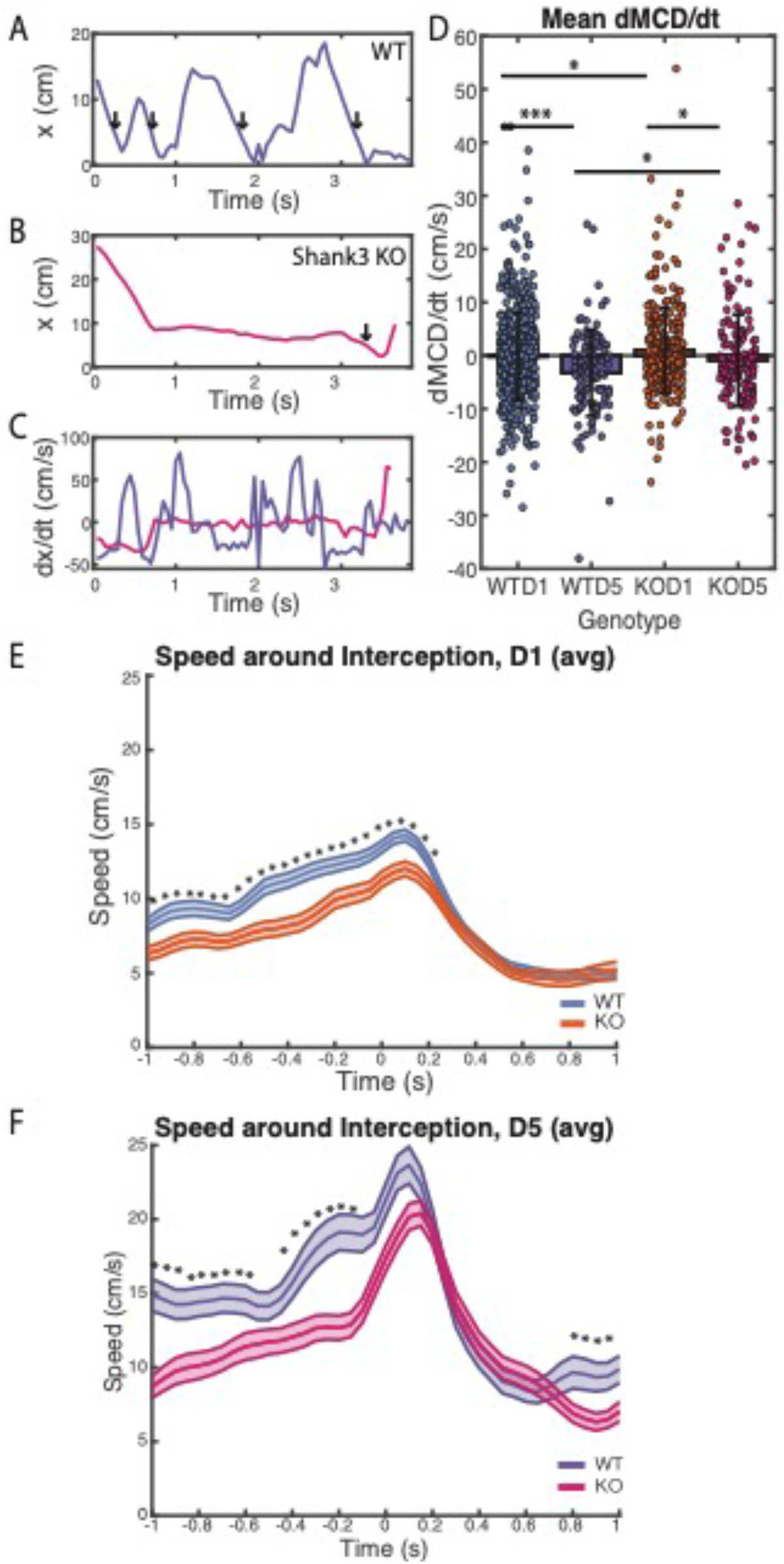
Shank3 KO animals exhibited similar average speeds as wild type animals during hunting approaches, but were slower to close in on crickets and exhibited less modulation of their speed immediately before interceptions. (A) 95^th^ percentile speed in the absence of appetitive stimulus, by session. 95^th^ percentile speed p=2.78e-4. Baseline speed was slower in Shank 3KO animals. (B) Average speed during approach, median over approaches during a trial. Wild type and Shank3 KO mice did not differ in average speed during approaches. Statistics generated through a repeated measures ANOVA: WTD1-KOD1 p=0.459, WTD5-KOD5 p=0.962, WTD1-WTD5 p=5.023e-04, KOD1-KOD5 p=6.477e-04. (C,D) Representative traces of the distance between mouse nose and cricket (mouse to cricket distance, x) during the course of one approach, for both wildtype (C) and Shank3-KO (D) mice after training. Vertical arrows indicate interceptions. (E) Derivative of mouse to cricket distance with respect to time, moving average over one second, for traces shown in (C) and (D). (F) Mean derivative of MCD with respect to time averaged over each approach period. Error bars indicate standard deviation. Calculated using two-sample t-test, WTD1-KOD1 p=0.048, WTD5-KOD5 p=0.026, WTD1-WTD5 p=2.29e-4, KOD1-KOD5 p=0.023. Experienced wild type mice exhibited lower values than experienced Shank3 KO mice, indicating that wild type animals closed the distance gap to the cricket more efficiently. (G-H) Mouse body speed one second before and after interception, averaged over all interceptions for a given genotype for both naïve (G) and experienced (H) mice. Shaded region indicates standard error. Asterisks indicate frames for which p<0.05 as calculated using a two-sample t-test. Wild type mice began to ramp up their speed about 500 ms before intercepting a cricket, while Shank3 KO mice did not greatly increase their speed until about 200 ms before intercepting a cricket.

## Results

To investigate whether juvenile mice are capable of learning to capture prey, and how this complex multimodal learning might differ in a mouse model of ASD, we compared littermate wildtype and Shank3 KO mice as they learned to hunt crickets. Our paradigm was adapted from that used successfully for adult mice (Hoy et al., 2016), but differed in that we first habituated animals to the hunting arena and exposed them to immobilized crickets. This was designed to reduce neophobia associated with approaching and consuming crickets, to allow us to focus on differences associated with the process of detecting, pursuing, and capturing prey.

### Juvenile mice readily learned to hunt crickets

The hunting behavior of juvenile mice was recorded with an overhead camera in a clear plexiglass arena of 14×14 inches (**Fig. 1A**). Before hunting, animals were habituated for two days, exposed to immobilized crickets, and were food-deprived the night before each session (**Fig. 1B**). Each trial began when a cricket was dispensed in one of four corners of the arena, and the behavior of both mouse and cricket was recorded on video. Sessions ended after the mouse had performed 5-6 trials (consuming 5-6 crickets), or after three hours, whichever came first. Animals were tested over five sessions, with one session occurring on each of five successive days.

Wild type juvenile mice hunting performance improved dramatically over these 5 hunting sessions. The fraction of crickets successfully captured increased (**Fig. 1D, E**), and the distance traveled before capture (**Fig. 1C, H**) and the time to capture (**Fig. 1I**) both decreased over the first few sessions, and reached asymptote by sessions 4-5. The average time to capture for these juvenile mice once they reached proficiency (∼15 seconds) was comparable to that reported previously for adult mice in a similar paradigm (Hoy et al., 2016). A video of an experienced wild type mouse hunting a cricket is shown in **Supplementary Video 1**.

### Shank3 KO mice learned more slowly and were less efficient at prey capture than wildtype littermates

There were dramatic differences between the two genotypes in their prey capture behavior. While most wild type mice successfully captured and consumed live crickets on the first day of hunting, most Shank3 KO animals did not, and they were slower to improve over successive sessions (**Fig. 1 D-I**).

To determine whether reduced motivation to consume a novel food source might contribute to the initial slowness to hunt, in a separate set of experiments we assessed the likelihood of mice to consume novel dead crickets or a novel cereal treat (Froot Loops). Shank3 KO mice were more variable than wildtype in the number of dead crickets consumed, and it took them longer to grasp crickets and initiate consumption (**Fig. 1J**); KO mice also consumed a smaller amount of cereal, although these differences in consumption were not statistically significant (**Fig 1F,G**). This suggests that Shank3 KO mice are more variable and slower than WT littermates in their willingness to grasp and consume a novel food source. Pre-exposure to immobilized crickets during habituation presumably reduced the effect of this neophobia on hunting performance.

Despite their initial slowness to hunt, over the course of 5 sessions Shank3 KO animals eventually became proficient at capturing live crickets, and reached a 100% success rate by the final session (**Fig. 1D, E**). A sample video of an experienced Shank3 KO mouse is shown in **Supplementary Video 2**. The rate of improvement was slower for KO than for WT littermates, and by session 5, KO animals traveled a longer distance (**Fig. 1H**) and took about twice as long to capture (**Fig. 1I**) as WT littermates (**Fig 1I**). Thus, juvenile Shank3 KO animals are able to learn a complex behavior requiring extensive multimodal sensorimotor integration, but did not become as efficient as their wildtype littermates.

### Naïve WT animals make more approaches during hunting than Shank3 mutants

To begin to understand what accounted for the differences in time to capture between genotypes, we carefully examined the instances where mice made an active “approach” to a cricket, define as periods of active pursuit. These approaches were manually scored for days 1 and 5 for both genotypes. “Interceptions” were defined as times when the distance between the mouse’s nose and the cricket’s body fell below a threshold of 5 cm. After an interception, it was possible that the cricket would be captured and eaten, or that the cricket would escape. If the mouse immediately continued to pursue a cricket after it escaped, then this was considered part of the same approach. Therefore, an approach might contain many interceptions. Approaches might end in a capture, or might end with a successful escape if the mouse did not continue an active pursuit after an unsuccessful interception.

On day 1 there were substantial differences in the number of active approaches by the two genotypes, with wild type mice making more approaches than KO mice (**Fig. 2A**). Additionally, on Day 1 both genotypes made many cricket interceptions (**Fig. 2B**). By day 5, the overall number of approaches and interceptions required to capture a cricket decreased for both genotypes (**Fig. 2A-B**) concomitant with an improvement in hunting performance.

The greater number of approaches of wild type mice on day 1 might reflect a higher intrinsic interest in crickets, or lower initial fear of crickets. Alternatively, the fewer approaches of Shank3 KO mice might reflect differences in sensory processing or motor planning. To examine these possibilities, we observed naive wild type and Shank3 KO mice as they interacted with novel dead crickets (**Fig. 2C**). The genotypes exhibited an equal propensity to investigate novel dead crickets, indicating that the differences in their interactions with live crickets is unlikely to result from differences in intrinsic interest or fear of novel objects. In order to understand whether a sensory deficit or a motor planning deficit might underlie the differences between the genotypes, we examined whether experienced mice would learn to begin their approaches to crickets from a greater distance. We assessed sensitivity to cricket presence by calculating the fraction of approaches starting with a Mouse-Cricket-Distance (MCD) greater than 15 cm (**Fig. 2D**). Both genotypes showed a numerical increase in this metric with training, suggesting that they became more able to recognize crickets as prey from a distance with experience. However, only wild type animals showed a statistically significant increase in the fraction of distant approaches.

### Shank3 KO mice exhibit increased periods of immobility compared to wild type mice

Although the two genotypes made similar numbers of approaches on day 5, the shorter time to capture and shorter path lengths traveled by wild type mice suggested that they were using time more efficiently than Shank3 KO mice. To begin to understand what contributes to these differences we next analyzed the time spent immobile during approaches (**Fig. 3A**) and found a striking difference: Shank3 KO mutants spent considerably more time immobile (movement <1cm/s) during approaches than their wild type littermates (WTD1 mean 0.42% time immobile, WTD5 0.039%, KOD1 0.99%, KOD5 0.26%). That is, while wild type mice smoothly pursued the crickets, the Shank3 KO mice exhibited a move-and-pause strategy during approach periods (see **Supplementary Videos 1-2**). WT mice showed an increase with training in the amount of distance travelled between one period of immobility and the next (**Fig. 3B**), and Shank3 KO animals learned to spend less time immobile overall (**Fig. 3A**). This indicates that both genotypes learned more active approach strategies over the course of the experiment.

Shank3 KO mice also spent considerably more time immobile in between bouts of cricket hunting, when they were free to explore the arena. In these periods when crickets were not present, Shank3 KO mice were immobile 39% of the time, compared to 27% of the time for wild type mice (**Fig. 3C**).

### Experienced Shank3 KO mice exhibit less modulation of speed during hunting than wild type mice

We next sought to understand whether the speed of the mice when moving differed between wild type and Shank3 KO mice. Speed was calculated from mouse body position assessed using DLC and averaged over two frames (Methods). “Moving speed” was taken to be the average speed calculated over manually scored approach periods.

When no cricket was present, Shank3 KO mice moved in the arena at slightly lower speeds than wild type mice (**Fig. 4A**). However, during live cricket approach, the average speeds of naive mice from both genotypes were identical, and both genotypes exhibited statistically significant increases in average approach speed as they became proficient hunters (**Fig. 4B**). Thus, when they did move during approaches, WT and Shank3 KO animals moved at similar average speeds.

We were curious to understand how the genotypes could exhibit identical average approach speeds on day 5, when wild type mice exhibited faster capture times than Shank3 KO mice. We found that wild type mice, in addition to spending less time immobile, were also more efficient when they did move. To quantify this, we calculated the average rate of change of the Mouse-to-Cricket Distance (dMCD/dt) during each approach, which is a measure of how quickly the mouse is closing on the cricket. Example MCD profiles during approach on day 5 are shown for wild type (**Fig. 4C**) and KO (**Fig. 4D**) mice, along with the corresponding profiles of dMCD/dt (**Fig. 4E**). Arrows in **Fig. 4C, D** correspond to interceptions. On day 1, mice of both genotypes exhibited highly inefficient movements, with a positive mean dMCD/dt in Shank3 KO animals indicating that the mice were less effective at approaching the cricket than the cricket was at escaping (WTD1 mean -0.164 cm/s, KOD1 mean 1.00 cm/s **Fig. 4F**). However, by day 5, both genotypes exhibited an overall negative dMCD/dt (WTD5 mean -3.26 cm/s, KOD5 mean - 0.905 cm/s), indicating that they had learned to close the distance to the cricket more effectively. Also on both days 1 and 5, wild type mice exhibited a more negative dMCD/dt than Shank3 KO mice. These results indicate that wild type mice were more efficient at closing the distance gap to the cricket compared to Shank3 KO animals.

In order to better understand how wild type mice closed the distance gaps more efficiently, we examined the speed of each mouse one second before and after an interception. All frames over this time range were averaged, for both naive (**Fig. 4G**) and experienced (**Fig. 4H**) mice of both genotypes. We found that wild type mice approached crickets more rapidly immediately before interception and that this difference was statistically significant for all frames during naive trials and for many frames during experienced trials. We additionally found that while experienced mice of both genotypes rapidly increased their speed as they were intercepting the cricket, wild type mice show an additional locomotive burst about 500 ms before interception. This suggests that wild type mice learned to modulate their speed more efficiently as they began to intercept a cricket.

## Discussion

We observed that both wild type and Shank3 KO juvenile mice can become proficient hunters, but exhibit a number of salient differences in learning rates and hunting strategy. Shank3 KO mice initially captured fewer crickets, and required more time and longer paths to do so (**Fig 1C-E, H-I**). After five days of hunting, smaller but significant differences in time to capture and path length between genotypes persisted. This slower time to capture was due to at least two factors: greater periods of immobility (pauses) during pursuit for Shank3 KO than wild type littermates; and because wild type mice learned to modulated their approach speeds to optimize capture in a manner that Shank3 KO mice did not. Thus, while both wild type and Shank3 KO mice are capable of learning this complex task, they differ in strategy and efficiency.

### Juvenile wild type mice can learn to hunt crickets

The developmental regulation of hunting behavior in rodents has not been systematically examined. A recent study comparing cricket hunting in very young mice (P20-22) and adult mice (Allen et al., 2022) found that young mice could learn to hunt crickets, although they did not approach crickets or artificial visual stimuli in a head-on manner to a degree that adults did, consistent with the idea that binocular visual fields in the visual cortex and superior colliculus are still immature. Here we show that by P28 (within the classical visual system critical period) juvenile mice are efficient learners, with learning curves that are quite similar to those reported for adult animals (Hoy et al., 2016; Galvin et al., 2021), with a strong improvement on the second day that continued through day 5. We also report here that, after learning, wild type juvenile mice modulate their behavior to accelerate rapidly toward the cricket and end with a burst of speed during the last 500 ms before interception.

### Shank3 KO and wild type juvenile mice show similar interest in crickets

In principle, the initial reluctance of Shank3 KO animals to hunt crickets could reflect lower interest in crickets, or greater short-term or long-term fear of crickets, compared to wild type animals. However, we did not find evidence for either of these potential explanations. During exposure to novel dead crickets, both genotypes showed a statistically identical number of interactions (**Fig. 2C**), which suggests that differences in performance cannot be attributed to a simple difference in interest or some type of neophobia that inhibits their examination of novel objects. This result is consistent with literature findings that wild type and Shank3 KO mice spent similar amounts of time exploring novel objects outside their nests (Wang et al., 2011).

Nevertheless, despite their interest in interacting with novel dead crickets, Shank3 KO animals consumed fewer of these crickets as compared to wild type mice (**Fig 1F**). Further, Shank3 KO mice consumed less novel fruity cereal than wild type mice (**Fig 1G**). These results suggests that Shank3 KO animals either a) exhibit a deficit in identifying a novel object as a potential food source, b) have a fear or other reluctance to actually ingest a novel food, and/or c) would like to eat the novel food but have difficulty carrying out the planning or the movements necessary to achieve the consumption. Any of these factors or a combination could have contributed to the lower performance of Shank3 KO mice on the first day of cricket hunting (**Fig 1C-E, H-I**). Regardless of naive KO behavior, statistical differences in the number of live crickets eaten (**Fig. 1D**) and probability of capture (**Fig. 1E**) disappeared after one to two days of hunting, suggesting that any such effects were transient.

### Wild type mice exhibit more dynamic hunting tactics

Shank3 KO and wild type mice exhibited substantial differences in pursuit movements during hunting, but these differences defy an easy characterization as a simple motor deficit. The two genotypes exhibited a pronounced difference in the time spent immobile – that is, briefly paused during ongoing behaviors such as approaches. Both genotypes showed a decrease in the amount of time spent immobile during approaches as a result of experience over the five days, but time immobile nearly disappeared for wildtype animals, while knockout mice continued to exhibit substantial pauses during pursuit. These pauses could reflect sensory and/or motor planning deficits, such that knockout animals need more time to process the sensory information about the cricket or to plan their next moves.

Wild type and knockout animals also exhibited substantial differences in their patterns of movement when they did move. Interestingly, the average speeds (**Fig 4B**) of both genotypes during approaches were not significantly different. However, over the course of an approach, wild type mice moved more efficiently, reducing their distance to the cricket more quickly than Shank3 KO mice (**Fig 4F**). The genotypes also exhibited major differences in how experienced mice modulated their speeds in the final second leading to an interception (**Fig 4H**). While Shank3 KO animals exhibited only one burst of acceleration as they reached the cricket, wild type mice moved faster 1 second before an interception and showed an additional increase in speed about 500 ms before interception. That is, experienced wild type mice employed an extra locomotive burst precisely when it was most needed: as they neared their peak speed, only centimeters away from their prey. This pattern was not present in wild type mice on the first day of hunting (**Fig. 4G**), so it represents a distinctive hunting tactic that wild type mice learned but knockout mice did not.

Shank3 KO mice exhibited lower levels of baseline ambulation when no cricket was present (**Fig. 3C**). This is consistent with previous reports, which show that Shank3 KO mice travel more slowly than their wild type counterparts during open field tests (Wang et al., 2011; Mei et al., 2016; Guo et al., 2019).

Therefore, while it appeared that Shank3 KO mice were capable of moving as quickly as wild type mice, they paused more, exhibited less efficient approaches, and did not modulate their speed in the same manner as wild type mice.

### Possible circuit mechanisms

Shank3 KO mice exhibited two major differences from wild type mice in cricket hunting. First, while Shank3 KO mice were interested in crickets, they appeared slow to recognize them as prey to be hunted and eaten (**Fig 1D-E, 2A-C**). Second, they paused more and were less efficient at modulating their speeds during approaches (**Fig 3,4**).

One explanation consistent with these data is that visual function is impaired by Shank3 loss. As homeostatic compensation and ocular dominance plasticity in the primary visual cortex is disrupted in Shank3 KO animals (Tatavarty et al., 2020), it is feasible that visual processing deficits contribute to lack of recognition of the cricket as prey as well as increased hesitation during hunting. This is consistent with the fact that vision (Hoy et al., 2016) and specifically binocular vision (Johnson et al., 2021) is required for effective cricket hunting, and is impaired in some monogenic mouse models with ASD features that exhibiting altered cortical homeostatic plasticity (Noutel et al., 2011; Durand et al., 2012; Felgerolle et al., 2019).

The superior colliculus has long been known to be involved in predatory hunting behavior as well as other goal-directed visual activities (Furigo et al., 2010; Wang et al., 2020). Global inactivation of the superior colliculus significantly degrades overall hunting performance (Zhen, 2017) while selective inactivation of defined cell types in the superior colliculus inhibits distinct aspects of prey approach and capture (Hoy et al., 2019), establishing that detailed circuit interactions mediated by this brain region have direct and well-defined effects on hunting behavior. Interestingly, the superior colliculus is involved in triggering the onset of predatory hunting (Shang et al., 2019; Huang et al., 2021), and Huang et al. found that inactivation of the pathway from the superior colliculus to the substantia nigra compacta disables hunting but does not disable exploratory behavior. It is possible that differences in superior colliculus or substantia nigra signaling or their connections might underlie some of the behavior differences reported here. While data is lacking in the literature as to the effect of Shank3 on SC function, Shank3 KO rats exhibit a lower relative volume of SC, suggesting that Shank3 loss may alter development of this brain region (Golden et al., 2021).

### Shank3 KO hunting behavior shows parallels with behavioral differences in autism-associated disorders

The genotype-dependent differences in both performing and learning cricket hunting observed in our experiments are analogous, in general terms, to some behavioral differences observed in autism-associated disorders. Firstly, Shank3 KO mice exhibited a greater amount of time immobile during hunting (**Fig. 3A**). Experienced Shank3 KO mice also pursued crickets less rapidly and efficiently (**Fig. 4F**). This is reminiscent of the behavior of individuals with ASD, who take more time to initiate goal-directed actions and move more slowly during the initiation of those actions (Glazebrook et al., 2006; Longuet et al., 2012; Alvares et al., 2016). Further, individuals with ASD exhibit irregularities in audiovisual integration (Stevenson et al., 2014; Brandwein et al., 2015; Feldman et al., 2018), and the availability of multisensory information (auditory and visual) improves cricket hunting in wild type mice (Hoy et al., 2016). This suggests that Shank3 contributes to effective learning of this complex ethological task in part by allowing mice to learn to rapidly integrate information about the position of their prey and respond to cricket movements expeditiously.

Another parallel between the behaviors observed in our study and symptoms of autism-associated disorders is that wild type animals exhibit a greater degree of improvement on multiple efficiency metrics than Shank3 KO animals. The lower learning proficiency observed according to all of these metrics is consistent with learning disability classically associated with Phelan-McDermid syndrome and other autism-associated disorders (Kolevzon et al., 2014), and furthermore generalizes known Shank3 KO deficits in operant conditioning (Bey et al., 2018), repetitive behaviors (Wang et al., 2016), and spatial learning and memory (Jaramillo et al., 2016; Jaramillo et al., 2017) to more complex and ethological behaviors.

## Acknowledgements

This work was funded by NIH EY022122 (SDV) and NIH EY025613 (GGT). We thank Derek Wise, Wei Wen, Brian Lane, Dan Leman, Hazal Uzunkaya, Francisco Mello.

